# Apoptotic proteins in *Leishmania donovani*: *In silico* screening, modelling, and validation by knock-out and gene expression analysis

**DOI:** 10.1101/2024.04.05.588228

**Authors:** Ketan Kumar, Lucien Crobu, Yvon Sterkers, Vijay Kumar Prajapati

## Abstract

Visceral leishmaniasis (VL), a life-threatening vector-borne illness that disproportionately affects children and elderly immunocompromised people, is a primary tropical neglected disease. No apoptotic partner proteins in *L. donovani* have been reported yet, which might contribute to the knowledge of parasite cell death and the establishment of alternative therapeutics. We used the Orthologues algorithm to search for the mammalian Bcl-2 family proteins orthologs, one anti-apoptotic and two pro-apoptotic, in *L. donovani*. We also included a pro-death aquaporin (AQP) protein due to its characteristic BH3 domain, which is known to interact with pro-apoptotic proteins in mammals. Molecular docking and molecular dynamics simulation studies were conducted to assess the protein-protein interaction between the identified apoptotic proteins and mimic mammalian intrinsic apoptotic pathways. The results showed that the pro-apoptotic protein interacted with the hydrophobic pocket of the anti-apoptotic ortholog, forming a stable complex, which may represent a critical event in the apoptotic pathways of leishmaniasis. To further establish an apoptotic pathway in *L. donovani*, we used several CRISPR-Cas9 approaches to target the identified proteins. The pure knocked population mutants, and episomal over-expressing mutant cells were exposed to apoptotic stimuli. TUNEL assay and quantitative expression profiling suggested that these proteins are needed during the parasite’s apoptosis and could play a role in the parasite’s survival.

**Author Summary:** Visceral leishmaniasis, a fatal systemic infection affecting internal organs, is one of three types of leishmaniasis in mammals alongside cutaneous and mucocutaneous leishmaniasis. It predominantly occurs in tropical and subtropical climatic zones, *Leishmania donovani* predominant in the Indian subcontinent and *Leishmania infantum* in the Mediterranean basin, the Middle East, Central Asia, South America, and Central America. This disease primarily affects children, immunocompromised adults, and the elderly. *L donovani,* transmitted by the infected sandflies complete its life cycle in humans, serving as reservoir. During its life cycle, at a particular stage, the parasite undergoes apoptotic-like events, yet underlying proteins or key factors remain unidentified. Using computational methods, we screened the *L. donovani* genome for potential candidate genes of the Bcl-2 family apoptotic proteins. We biologically/experimentally validated our *in-silico* findings using molecular editing tools like CRISPR-Cas9, advancing our understanding of the parasite’s apoptotic pathway. Targeting this pathway could lead to more effective therapeutics against visceral leishmaniasis.

## Introduction

Visceral leishmaniasis (VL) is a vector-borne chronic disease in children and elderly immunocompromised people. It is still a major tropical and/or sub-tropical neglected illness. Its prevalence is influenced by sandfly species traits, local ecosystem dynamics of transmission locations, current and prior parasite exposure to the human population, and human behaviour (1). Since VL was first studied in India over a century ago, researchers have seen incidence cycles that climb and fall with some regularity. According to WHO, an estimated 50000 to 90000 new cases of VL are diagnosed each year across the world, of which 90% of new cases are recorded in three eco-epidemiological hotspots regions, including the following ten countries: Brazil, China, Ethiopia, Eritrea, India, Kenya, Somalia, South Sudan, Sudan, and Yemen (2). In the Indian subcontinent, Humans are the principal reservoir for VL. The causative agent of VL is a protozoan parasite *Leishmania donovani* in the Indian subcontinent and transmitted by sand fly, Phlebotomus argentipes (3). At the same time, VL is caused by *Leishmania infantum* in the Mediterranean basin, the Middle East, Central Asia, South America, and Central America (4).

A number of chemotherapeutic options are available to control the spread of VL in the endemic zones. Chemotherapeutic (5) agents like pentavalent antimony, amphotericin B, miltefosine, paromomycin, pentamidine, and liposomal amphotericin B (6) are used over time with their advantages and disadvantages. Immunotherapy is a relatively new approach to modulating the host’s immune response with effective biological molecules. Even though immunotherapy constitutes a wide range of therapeutics and prophylactics, this approach is in its early stage of development (7). These currently available treatments elicit undesirable side effects, show administration limitations, and costlier therapeutic interventions, and the advent of drug-resistant genotypes render this approach ineffective (8). These limitations in leishmaniasis are a severe worldwide health concern, especially in developing and endemic countries.

To address these constraints, efforts are constantly made to enhance understanding of the aetiology and pathogenesis of leishmaniasis (9). In this context, it is interesting to cite that leishmanial parasites undergo regulated apoptotic-like cell death during their course of life cycle and pathogenesis, phenotypically similar to higher eukaryotic apoptosis (10). *Leishmania* maintains the clonality inside the sandfly gut by managing the promastigotes population, choosing infectious forms ideal for disease transmission. Promastigotes that do not mature into virulent metacyclic forms are terminated in a regulated manner to help the metacyclic by restricting the limited nutrition supply of the procyclic (11). Parasite density within the host is also controlled to prevent hyper parasitism, host mortality, and the inhibition of parasite transmission (12). In addition, macrophage phagocytizing apoptotic *Leishmania* cells inhibit pro-inflammatory cytokines secretion and increase the anti-inflammatory chemicals (13,14). One can infer that *Leishmania* has retained such apoptosis-like cell death mechanisms during evolution because they are advantageous or necessary for the survival of species or populations (14,15). In contrast to higher eukaryotes, understanding apoptosis-like cell death mechanisms in *Leishmania* is still inadequate. This is because the molecular machinery required for initiating and executing apoptosis-like cell death is yet to be identified and defined (16). Furthermore, it is crucial to understand protein-protein interactions that necessitate cellular structure, function, and organisation (17). Protein-protein interactions mediate most biological activities; hence, a complete description of the association process at the molecular level is required to understand the fundamental mechanisms. Molecular docking provides a tool for basic research by computationally predicting the interaction of two molecules. Protein-protein docking techniques, for example, seek to discover the natural binding mechanism between two proteins. In this study, we did the cross-search for orthologous genes through the genome of *Leishmania donovani* for mammalian essential apoptotic proteins, Bcl-2 (18). Bcl-2 family proteins control and cause mitochondrial outer membrane permeabilization, an essential process in apoptosis and an ideal candidate for orthologous gene search. Orthologous genes are one of the two fundamental homologous genes that arose through speciation from a single ancestral gene, hence representing the evolution of species (19). All Bcl-2 family proteins have at least one conserved Bcl-2 homology (BH) domain that is required for the significant apoptotic function and interactions. Based on its primary function, the Bcl-2 family is divided into three groups: (a) anti-apoptotic proteins, (b) pro-apoptotic pore-formers, and (c) pro-apoptotic BH3-only proteins (20). We found orthologs of anti-apoptotic and pro-apoptotic proteins and docked them. *L. donovani*’s BH3 domain-containing protein was likewise chosen and docked with anti-apoptotic orthologs. The stability of the docked complex was assessed using a molecular dynamics simulation study, which revealed a plausible apoptotic route involving an anti-apoptotic protein and a protein with a BH3 domain. In a second stage, we wanted to use reverse genetics to obtain experimental validation and demonstrate the role of the proteins identified by the bioinformatics analysis in the apoptotic process. Thus, overexpression of tagged proteins and CRISPR-Cas9-based edition were used. The generated mutant parasites were then characterised by quantitative expression profiling in the presence and absence of an apoptotic stimulus. Our *in silico* and experimental findings suggest, for the first time, the significance of the identified protein in apoptosis and propose a putative interacting model for *Leishmania* apoptosis or apoptosis-like event.

## Methods

### Search for apoptotic ortholog of L. donovani

Orthologous gene finding is significant in comparative genomics across all disciplines of biology. Identification of orthologs between species, for example, might yield functional insights for genes with unknown functions and is an essential step in evolutionary reasoning (21). Most ortholog identification techniques currently use all BLAST comparisons, which in itself is time and memory expensive. Here, in-house OrthoMCL (22) from eukaryotic pathogen, vector, and Host informatics resources (VEuPathDB) was employed to identify orthologs using the Markov cluster approach (23). OrthoMCL is a large-scale genome-wide scale algorithm that begins with complementary best BLAST hits within each genome as putative ortholog pairs and then aligns relevant proteins in a similarity network. Afterwards, MCL (Markov clustering algorithm) mega clustering tool differentiates each protein pair considering evolutionary distances (24).

### Structure modelling of the resultant apoptotic ortholog protein

The orthologs search results subsequently did not have the three-dimensional structure resolved to date. As a result, structure and structure details about the protein’s binding site and binding residues are unavailable. Hence, for the *in-silico* structural prediction of protein, the sequence of orthologous explored within the genome of *L. donovani’s* BPK strain in reference to mammals was retrieved from the VEuPathDB (https://veupathdb.org/veupathdb/app, last access March, 18, 2024). The protein sequences were used for the 3D structure prediction by exploring the Phyre2 web-based server to predict protein structure (25).

### Structure refinement and validation of modelled proteins

The modern template-based modelling approaches fail to produce realistic structures for less-conserved local regions even when the overall structure can be reliably predicted. Less-conserved or unreliable local regions (ULRs) are frequently associated with functional specificity, and ab-initio refinement is invaluable for further functional and design studies (26). So, the predicted structures were further refined on GalaxyWeb refine web-based server to attain its nearest native 3D conformation (27). Furthermore, the quality of refined models was validated by analysing QMEANDisCo (28) score and their structural conformations on the Ramachandran plot using the structure assessment tool of the SIWSS-MODEL server (https://swissmodel.expasy.org/assess, last access March, 18, 2024) (29).

### Molecular docking to investigate potential protein-protein interactions

Molecular docking is a computational tool that generates critical structural information and interaction between protein-protein and/or protein-ligand molecules. The molecular docking algorithm ranks and picks potential hits based on their binding energy score (30). ClusPro, HDOCK, and pyDock online docking packages were employed for docking investigations. ClusPro server is a widely used protein-protein docking server that uses PIPER, a docking tool, to execute a protein-protein docking study (31). The docked complexes are ranked using the PIPER scoring system, and the information from SAXS is utilized to filter the produced structures. This methodology enables proteins to be docked without prior knowledge of the complex’s structure, yielding models defined by dense clusters of low energy docked items at their centres. The HDOCK server is a macromolecular docking suite with a high level of integration for reliable and quick protein-protein docking (32). It employs a hybrid docking approach that integrates empirical knowledge about the protein-protein binding site and SAXS during the docking and post-docking operations. Among docking servers, the HDOCK server is unique in that it predicts the interaction interface between two interacting proteins. Its efficient integration of biological and experimental data aid in investigating the de-novo protein complex. pyDockWEB is a rigid-body docking webserver that provides non-expert users with quick access to state-of-the-art docking prediction in a five-step procedure (33). pyDockWEB uses a new custom parallel FTDock implementation to produce docking poses that can scale to many processing cores while maintaining the same prediction accuracy. FTDock scoring creates an efficient empirical potential, consisting of electrostatics and desolvation terms with a little Van der waals energy contribution. Consequently, this webserver returns the best ten docking orientations.

### Molecular Dynamics Simulation for the validation of protein-protein interaction

Molecular dynamics simulation is a popular tool to analyse the characteristics of structures and their microscopic interactions (34). Molecular dynamics simulations are a powerful, advanced tool to understand molecular processes at a resolution that is often unattainable by experiment and explore atomic-level mechanisms that are often outside the purview of existing experiment methods (35). Molecular dynamics simulations and other variations are rapidly being employed to investigate more complicated biological systems from the electron to the molecule level. Stable conformations, folding, energy landscape, transitions between macro-states, and aggregation states might be investigated using Molecular dynamics simulations and other simulation approaches (36). Organic compounds, small peptides, protein-protein interactions, lipid-protein complexes, and even viral capsids have been molecularly modelled using molecular dynamics. Here, UAMS-Simlab WebGro Macromolecular simulation was used for Molecular dynamics simulation. WebGro is a fully automated and easy-to-use computational biology tool for all researchers (WebGRO for Macromolecular Simulations; https://simlab.uams.edu/, last access March 18, 2024). For completely solvated molecular dynamics simulations, WebGro employs the GROMACS simulation program (37). Here, Molecular dynamics simulations were executed to investigate the stability of protein-protein complexes in an explicit water solution at 300K temperature. The protein complex combinations were placed in a triclinic box with a GROMOS96 43a1 force-field. By introducing NaCl counter ion, the solvated system was neutralized. Solutes were treated to NVT/NPT ensemble to equilibrate the system.

### Parasite culture

Promastigotes form of L donovani strain 1S2D (MHOM/SD/62/1S-CL2D, referred to here as LD1S) was grown at 27°C, in chemically defined HOMEN culture with 10% FCS and 0.005% Hemin (3.5 mg/mL) (38). The cell culture density at the start was 1×10^6^ cells/mL, as assessed by the CellDrop FL (DeNovix automated cell counter).

### Generation of Cas9 expressing parasites

Plasmid pTB007 (39) for T7 RNA polymerase and Cas9 expression was transfected into LD1S to achieve LD1S-T7-Cas9-expressing cells. The plasmid also conferred hygromycin resistance, which was used to select transfected cells. In the Amaxa Nucelofactor IIb (Lonza) with 1X Tb-BSF transfection buffer, one pulse with program X-001 was used for transfection.

### Primer design for Cas9 knock-out & quantitative PCR

The primers for amplifying donor DNA and sgRNA targeting genes of interest (GOIs) were created as described in Beneke et al (39). Briefly, we used the Primer design tool on the LeishGEdit (http://leishgedit.net/) website. The target sequences of the GOIs were obtained from TriTrypDB (https://tritrypdb.org/tritrypdb/app, last access March 18, 2024) using their unique gene ID. To ensure specificity, the primer sequences were checked against to the complete genome using TriTrypDB BLAST tool. Donor DNA with Puromycin and Gentamicin drug resistance was amplified by PCR using the pTPuro and pTNeo plasmid backbones. The quantitative PCR primers were designed using Roche’s LC480 probe design software, which the quantitative PCR manufacturer provided. The primer design used SYBR green with a Tm target 60 and a size product smaller than 200 as parameters. IDT Belgium was contacted for the commercial synthesis of all designed primers (see supplementary list).

### Generation of mutant parasites

LD1ST7-Cas9-cells were transfected with PCR-amplified donor DNA for KO and sgRNA template targeting the GOI. The concentration used per transfection of donor DNA and sgRNA used were in the range of 25-30 µg (2 KO cassettes and 2 sgRNA templates). For overexpression and addback (AB) experiments, the GOI was cloned into a pTH6nGFPc vector (40) in frame with a GFP tag. The resulting plasmid was transfected into LD1S promastigotes. For all transfections, cells were in exponential phase with cells in the range of 5-6 x10^6^ cells/mL.

### Apoptotic induction of mutants

Miltefosine (MLT) stock of 20 mM was prepared in dH_2_O (SIGMA M5571-50 mg) and was used as apoptotic stimuli for induction of apoptosis of cell. KO and AB mutants prepared were treated with 5 µM and 10 µM MLT (IC_50_ of MLT = 13.2 ± 0.8 µM) concentration for 24 h. Growth phase cell of mutants was used for MLT treatment. Post 24 h MLT treatment 1 mL of cells were used for the TUNEL assay for apoptotic detection and rest were used for the RNA extraction and reverse transcription for qRT-PCRs.

### TUNEL assay

*In-situ* detection of DNA fragments following treatment of cells with MLT 5 μM and 10 μM for 24 h was measured by TUNEL using a Cell Death Detection kit (In-Situ Cell Death Detection Kit, POD; Roche; Cat. No. 11 684 817 910). 1000 μL of cells were centrifuged (3000 rpm for 5 min), resuspended in 1000 μL PBS 1Xwere, washed with PBS 1X, and fixed with 500 μL of 4 % paraformaldehyde for 1 h. 40 μL of fixed cells were deposed on ThermoFisher Scientific Menzel-Glaser Superfrost Plus Gold slide per well and air dried. Wells were washed with PBS 1X and incubated with H_2_O_2_ (3 % in methanol) for 10 min at 4°C. Then washed twice with PBS, placed on ice and permeabilized with freshly prepared, chilled 0.1 % sodium citrate in 0.1 % Triton X-100 solution for 2 min. For labelling preparation, the positive control well was treated with DNAse (DNase I, EN0525 – 1 U/L; Invitrogen) for double-strand break. A negative control cell was added with 50μL of Vial 2 of TUNEL kit without terminal transferase, and 50 μL of Tunel mixture containing terminal transferase was added per well, including the positive control well. Slides were incubated in a humidified chamber at 37°C for 1 h and washed with PBS. Cells were then incubated with 30 μL of diluted Hoechst 33342 (Thermo Fisher Scientific, # H3570) for 5 min and rewashed with PBS 1X, and 7 μL of antifade reagent (SlowFade^TM^ Gold, Invitrogen ref S36937) was added in each well. Slides were covered with a 1.5H cover slip.

### Mitotracker staining and fluorescence

Mid-log LdAQP-GFP cells at a density of approximately 3×10^6^ cells per mL were incubated with Mitotracker (Invitrogen/Molecular Probes, #M7512) at a final concentration of 40 nM during 15 min at 27°C in culture medium without FCS. Culture was then washed three times and allowed for one additional hour of growth in complete medium. Cells were washed twice in PBS1X. Nuclear and kinetoplast DNAs were stained with Hoechst 33342 (Thermo Fisher Scientific, # H3570 and antifading was added in the mixture. A small volume of concentrated cells was deposited on poly-lysin home-coated slide. Slide was mounted with a 1.5H coverslip.

### Image acquisitions

All images were acquired using Zen software, on a Zeiss® Axioimager Z1 microscope equipped with ORCA-Flash 4 OLT CMOS camera (Hamamatsu®) with 63x objective (Plan-Apochromat 63x/1.40 Oil) and filters adapted for the different fluorescence used.

### RNA extraction and cDNA synthesis by reverse transcription

MLT post 24 h treatment cells were recovered by centrifugation at 3000 rpm for 3 minutes, and total RNA was extracted using the RNeasy Mini Kit protocol (Qiagen-ref 74104). Extracted total RNA was treated with Turbo DNAse (Invitrogen^TM^ Ref: AM1907) to achieve the perfect genomic DNA elimination. Post DNAse treatment, the extracted RNA was transcribed into cDNA using a kit (Invitrogen^TM^ SuperScript^TM^ III First-strand Synthesis Super Mix for qRT-PCR, Ref. 11752050) containing Oligo dT with PolyA tails, enabling the reverse transcription of mRNAs. No-RT treatment was also used for the qPCR as a negative control.

### Primer pair and qPCR efficiency

Primer pairs for quantitative PCR were assessed on genomic DNA for PCR efficiency, reproducibility, and primer pair efficiency. Seven dilutions (D) of gDNA starting 1 ng/µL concentration as D1 with 10-fold decreased concentration till D7 were used for the primer pair efficiency and performed in triplicates for each primer pair mix. Primer pair mix with 7 µL of 2X SYBR green (LightCycler 480 SYBR Green I Master; Roche; Cat No. 04707516001) and 1 µL of 10 µM of forward and reverse primer were prepared for all the gene targets, including the reference gene. 5 µL of gDNA dilutions were added in triplicate, and 5 µL H_2_O as a negative control in triplicate was included on 96-well plates. The PCRs were performed on Roche Light Cycler 480 with run specifications: Tm = 60℃, elongation time = 40 seconds with 45 cycles.

### Quantitative expression of GOIs in wild type and mutants

cDNA prepared from mRNA by First-strand synthesis Superscript III reverse transcriptase were amplified in Roche LightCycler 480 Real-Time PCR machine using 2X SYBR green. The PCR conditions were 95°C denaturation, followed by a combined annealing and elongation step at 58°C. The data were analysed using the LightCycler 480 software against a standard curve obtained from serial gDNA dilutions. cDNA concentration of 100ng/µL was used in triplicates with water and no-RT treated RNA as negative control. Data from three independent experiments for wild type and mutant cells were normalized to the cGAPDH (LdBPK_362480.1) as a reference gene.

## Results

### In silico screen for identifying L. donovani proteins involved in an apoptotic pathway

#### Bcl-2 family proteins in L. donovani

The Bcl-2 family protein comprises both pro and antiapoptotic proteins. One ortholog for antiapoptotic findings, LdBPK_302600.1, against antiapoptotic BCL-2 and two pro-apoptotic ortholog search results, LdBPK_291360.1 & LdBPK_020680.1, against Pro-apoptotic Bcl-2 family proteins were obtained in L. donovani’s BPK strain using the OrthoMCL algorithm. TriTrypDB registered Gene Id LdBPK_302600.1 with potential RNA-binding protein function and gene IDs LdBPK_291360.1 and LdBPK_020680.1 with putative HSP78 and ATP-dependent Clp protease subunit functions and as a conserved hypothetical protein. A pro-death BH3 domain containing putative AQP protein, gene Id LdBPK_221270.1, was also selected for its putative pro-apoptotic activity because of the presence of the BH3 domain and its possible interaction with anti-apoptotic orthologs (41). The BH3 domain is required for the BCL-2 family member’s primary apoptotic function and intracellular membrane interactions (42).

#### Homology modelling of apoptotic proteins

Protein structures were constructed using the homology modelling technique because they were unavailable in the protein data bank. The homology model of identified Bcl-2 family proteins and BH3 domain-containing AQP protein of *L. donovani* was produced using the Phyre2 homology modelling server for 3D structural characterization. Phyre2 modelled the proteins with 78 percent residue modelled with more than 90% confidence for LdBPK_302600.1 (AAPO), 99 percent residue modelled with 90% confidence for LdBPK_291360.1 (PAPO-I) and 88 percent residue modelled with 90% confidence for LdBPK_020680.1 (PAPO-II) and LdBPK_221270.1 (AQP). AAPO and PAPO are acronyms used hereafter throughout the manuscript based on the Anti/Pro apoptotic putative role based on our *in silico* finding). The modelled protein was refined using the GalaxyWeb refine webserver and validated for final structure using the SWISS-MODEL structural assessment tool. According to the Ramachandran plot investigation, the most preferred and allowed locations for the statistical distribution of ϕ and ψ angles combination of amino acid residues were 91.92% for AAPO, 91.2% for PAPO-I, 92.27% for PAPO-II and 93.66% for AQP, indicates the high accuracy of the models (Fig. 1A-B).

**Fig. 1:**
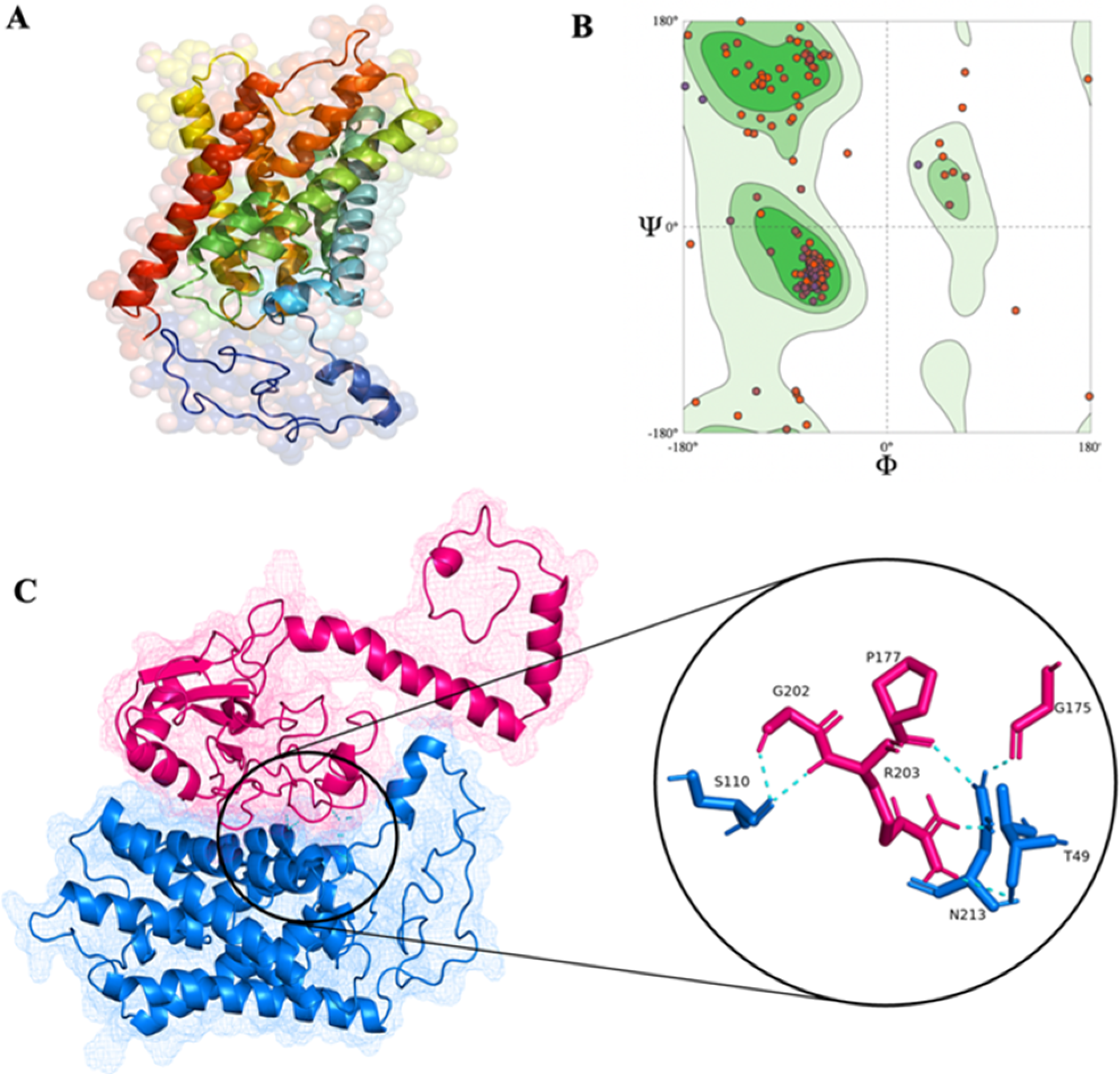
In silico AQP structure and binding prediction. (A) Homology structure modelled by Phyre2 for AQP; (B) Ramachandran plot for ϕ and ψ angles for the validation of homology predicted AQP structure; (C) Docking prediction of AAPO in pink and AQP in blue with their interacting residue in the inset image

### Molecular Docking studies for the interaction of Anti-apoptotic with Pro-apoptotic proteins

The main objective of this work was to scan apoptotic proteins and their interactions in *L. donovani* to see if there were any plausible pathways for the evolutionary conserved phenomena of apoptosis. Pro-apoptotic proteins and BH3 domain-containing proteins were docked with an anti-apoptotic protein in three combinations. Firstly, PAPO-I was docked with AAPO; secondly, PAPO-II with AAPO; and thirdly, AQP with AAPO. Pro-apoptotic proteins served as receptors for the docking experiment, and anti-apoptotic proteins served as ligands. These molecular docking studies were carried out using three protein-protein docking servers, each with a different docking algorithm. Out of the three docking platforms, the ClusPro inbuilt algorithm of energy minimization refining of chosen structures aids in selecting its docked complex for the subsequent molecular dynamic simulation research stage. ClusPro docked complex resulted in the best cluster complex due to its PIPER-based docking and RMSD-based grouping of lowest-energy complexes to identify one of the most relevant clusters that reflect the most credible complex models.

Identification of the Pro-apoptotic binding sites on Bcl-2 anti-apoptotic protein of L. donovani

Pro-apoptotic and anti-apoptotic proteins were randomly docked in two combinations, PAPO-I + AAPO (combination I) and PAPO-II + AAPO (combination II) (Suppl Fig. 1). It was found that the docked score trends between these two combinations were comparably similar except for HDOCK (Table 1). The alignment and polar interaction residues for AAPO were similar in both combinations, with a higher docking score for combination II. Combination II has a better docking score, indicating a believable complex. The PAPO proteins are 42.22% similar by sequence similarity.

**Table 1:**
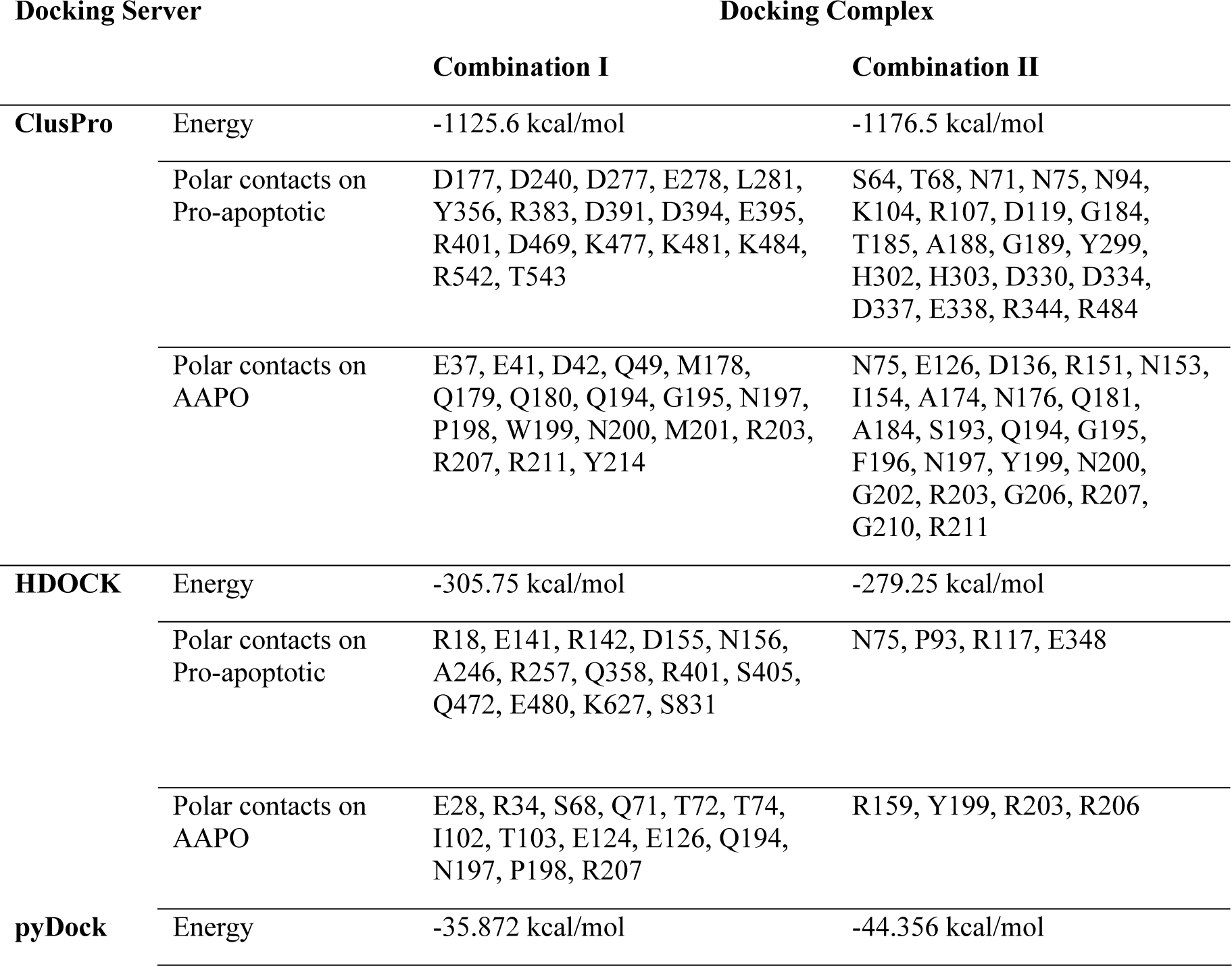
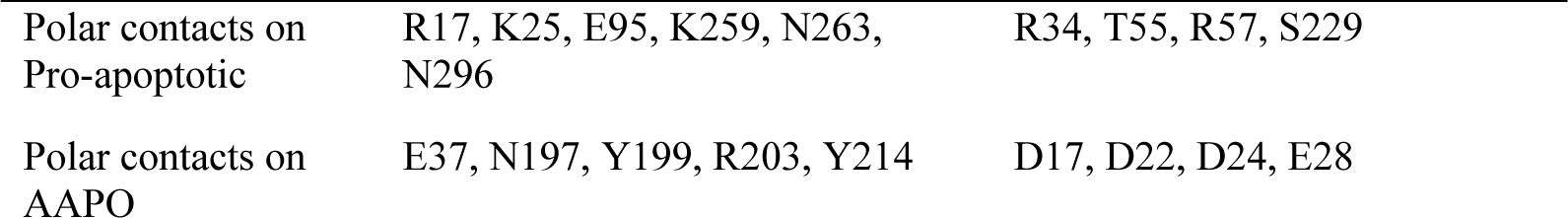
Docking energies of Combination I (PAPO-I + AAPO) & II (PAPO-II + AAPO) with their interacting residues in kcal/mol

| Docking Server |  | Docking Complex |  |
| --- | --- | --- | --- |
|  |  | Combination I | Combination II |
| ClusPro | Energy | -1125.6 kcal/mol | -1176.5 kcal/mol |
|  | Polar contacts on Pro-apoptotic | D177, D240, D277, E278, L281, Y356, R383, D391, D394, E395, R401, D469, K477, K481, K484, R542, T543 | S64, T68, N71, N75, N94, K104, R107, D119, G184, T185, A188, G189, Y299, H302, H303, D330, D334, D337, E338, R344, R484 |
|  | Polar contacts on AAPO | E37, E41, D42, Q49, M178, Q179, Q180, Q194, G195, N197, P198, W199, N200, M201, R203, R207, R211, Y214 | N75, E126, D136, R151, N153, I154, A174, N176, Q181, A184, S193, Q194, G195, F196, N197, Y199, N200, G202, R203, G206, R207, G210, R211 |
| HDock | Energy | -305.75 kcal/mol | -279.25 kcal/mol |
|  | Polar contacts on Pro-apoptotic | R18, E141, R142, D155, N156, A246, R257, Q358, R401, S405, Q472, E480, K627, S831 | N75, P93, R117, E348 |
|  | Polar contacts on AAPO | E28, R34, S68, Q71, T72, T74, I102, T103, E124, E126, Q194, N197, P198, R207 | R159, Y199, R203, R206 |
| pyDock | Energy | -35.872 kcal/mol | -44.356 kcal/mol |

|  |  |  |
| --- | --- | --- |
| Polar contacts on Pro-apoptotic | R17, K25, E95, K259, N263, N296 | R34, T55, R57, S229 |
| Polar contacts on AAPO | E37, N197, Y199, R203, Y214 | D17, D22, D24, E28 |

### Identification of AQP binding sites on Bcl-2 anti-apoptotic protein of L. donovani

In contrast to the previous combinations, AQP + AAPO (combination III) (Fig. 1C) showed superior binding energy (Table 2). In combination III, hydrophobic amino acid residues were involved in AAPO interaction with AQP, and their orientation in the docked complex is through the hydrophobic surface.

**Table 2:** Docking energy of combination III (AQP + AAPO) with their interacting residue in

| Server | Docking Complex | Combination III |
| --- | --- | --- |
| **ClusPro** | Energy | -1448.9 kcal/mol |
|  | Polar contacts on AQP | T49, S110, N213 |
|  | Polar contacts on AAPO | G175, P177, G202, R203 |
| **HDOCK** | Energy | -61.634 kcal/mol |
|  | Polar contacts on AQP | F102, S106, S110, N213 |
|  | Polar contacts on AAPO | Q182, I182, G202, R203 |
| **pyDock** | Energy | -317 kcal/mol |
|  | Polar contacts on AQP | S99, N213 |
|  | Polar contacts on AAPO | N197, R211 |

Anti-apoptotic proteins in mammalian intrinsic pathways adopt a highly conserved tertiary structure, forming a hydrophobic BH3 domain-binding grove that behaves as a binding site for other BH3 domain proteins (43,44). The BH3 domain is required for primary apoptotic function and interplay with intracellular membrane proteins (45). We did not find binding through the BH3 domain of AQP in mammals. We did, however, locate hydrophobic amino acids at AAPO protein-interacting sites. Combination III was the most successful, forming a hydrophobic domain groove through which the interaction occurs.

### Molecular dynamics simulation studies of docked complex suggest a model for the stable interaction

We conducted a molecular dynamics simulation study that aims to understand the conformation changes and evaluate dynamic behaviour of protein-protein complex for consistency of interaction (46). Finally, Prism software was used to generate root-mean square deviations (RMSD), Root Mean Square Fluctuation (RMSF), radius of gyration, and solvent-accessible surface area dynamics trajectory graphs to compare the protein complex’s behaviour throughout the simulation (47).

### Assessing protein stability and equilibrium dynamics through molecular dynamics simulation: A comparative RMSD analysis

During molecular dynamics simulation, the combinations RMSD are determined by translating and rotating the Cα atom coordinates of the instantaneous structure to superimpose with the reference structure, i.e., the initial beginning structure with the most significant overlap. Thus, RMSD estimates the deviation of the structures from their overlap and utilizes it to validate the model whether the systems are in equilibrium and do not encounter any conformational changes. A suitable solvent model should produce minimal RMSD values that become consistent throughout the journey. To evaluate the flexibility of the protein’s backbones, the RMSD was compared for the three complex system combinations and was observed to be less than 1 nanometre (nm) over the simulation period. Combinations I and II had average RMSDs of 0.98 and 0.81 nm, respectively, and combination III had an RMSD of 0.67 nm, showing that all complexes are coherent. As depicted (Suppl. Fig. 2A), the overall equilibrium for combination II was attained at 15 ns with an RMSD of 0.08 nm, whereas the equilibrium for combination III was at 5 ns with an RMSD of 0.06 nm. In the last 40^th^ ns of the simulation, a plateau graph was observed, which becomes stable shortly after 5ns and continues below 1 nm for combination III, indicating that perhaps the system is steady and well equilibrated. Except for combination I, the RMSD computed for combinations II, and III revealed no significant variations in the backbone, showing that the protein binding is stable and robust, with minimal conformational changes that do not affect the protein backbone stability during the simulation.

### Exploring amino acid dynamics: root-mean square fluctuation (RMSF) analysis reveals protein-protein interaction dynamics

We explored the RMSF fluctuations of simulation studies to monitor the dynamic behaviour of amino acid residues during protein-protein interactions (48). RMSF measures the average divergence of amino acid residues and computes individual residue flexibility, that is fluctuation of residues throughout the dynamics from the initial structure. As a result, RMSF examines the parts of the structure that deviate the greatest from their mean structure (or least). RMSF per residue is often plotted versus residue number and can suggest which amino acids in a protein contribute the most to molecular motion biologically. The degree of chain flexibility in the three combinations was represented by the RMSF of each residue from its average time location (Suppl. Fig. 2B). Internal fluctuations, as well as conventional N-and C-terminal fluctuations, were found in all apoptotic proteins of *L. donovani*. Aside from the N-and C-terminal regions, the flexible area in the protein region, includes the residues that belong to the site linked explicitly to its protein partner interaction sites, as shown in Tables 1 and 2. Several other flexible residues were detected along with these, that are thought to be allosteric protein sites because higher fluctuations or structural alterations occur in regions known to be critical for binding and/or catalysis.

### Assessing protein compactness through radius of gyration fluctuations

We plotted the radius of gyration (Rg) for different time frames in a dynamic system to evaluate the compactness of the structures as an indicator of the size and compactness of all combinations (49). Over the dynamics period, a stably folded protein maintains a reasonably constant Rg. Based on the Rg plot (Suppl. Fig. 2C), we can conclude that among the three combinations, combination III had the lowest average Rg of 2.36 nm, while combinations I and II had average Rg of 3.72 and 3.52 nm, respectively. Combination III had a relatively steady Rg after the 7^th^ ns over the period of dynamics. On the other hand, Rg of combination I and II are rocketed and fluctuated. The evident lowest and slightest variation Rg for combination III indicates that the protein is stable and compact, with the tightest protein packing.

### Assessing solvent accessibility and stability in protein complexes

We plotted the SASA (Solvent Accessible Surface Area) of protein complexes to evaluate the accessible surface area of the complexes accessible to solvent. SASA is the surface area of a protein that evaluates the interaction of complexes with their solvent molecules (50). During a 50 ns Molecular dynamics simulation, average SASA values for the combination I, II, and III were recorded (Suppl. Fig. 2D) to be 503.46 nm^2^, 458.55 nm^2^, and 207.17 nm^2^, respectively. SASA values for combination I was noticeably decreasing with fluctuation during the simulation period; however, the SASA values of combination II & III did not fluctuate much and decreased over time, indicating structural compactness (51).

The trajectory graphs of the apoptotic protein complex’s behaviour during the molecular dynamic simulation indicates that the protein-protein interaction of AAPO and AQP is exceedingly stable than the other two combinations. The result revealed that the protein backbone of AQP with AAPO remained stable throughout the simulation period, with no notable fluctuations. AQP interacts with the hydrophobic surface AAPO to form a complex with a low binding score similar to mammalian anti and pro-apoptotic interactions. The backbone of LdPAPO-II, on the other hand, showed slight anomalous variations throughout the simulation period, whereas LdPAPO-I demonstrated instability with the AAPO.

### Experimental validation of in silico hits

#### Exploring the impact of AQP gene expression on Leishmania apoptosis-like events

The first protein we targeted was AQP to check the effect of a characteristic BH3-domain containing the LdAQP gene on parasite cell death or apoptosis-like events. We successfully performed a Cas9 KO (Fig. 2A), a complementation (add-back) of the KO and an episomal overexpression in the wild type. This enabled us to look at the localisation of the protein, the changes in expression of the other target proteins as a function of AQP expression and the percentage of TUNEL-positive cells under these different conditions.

**Fig. 2:**
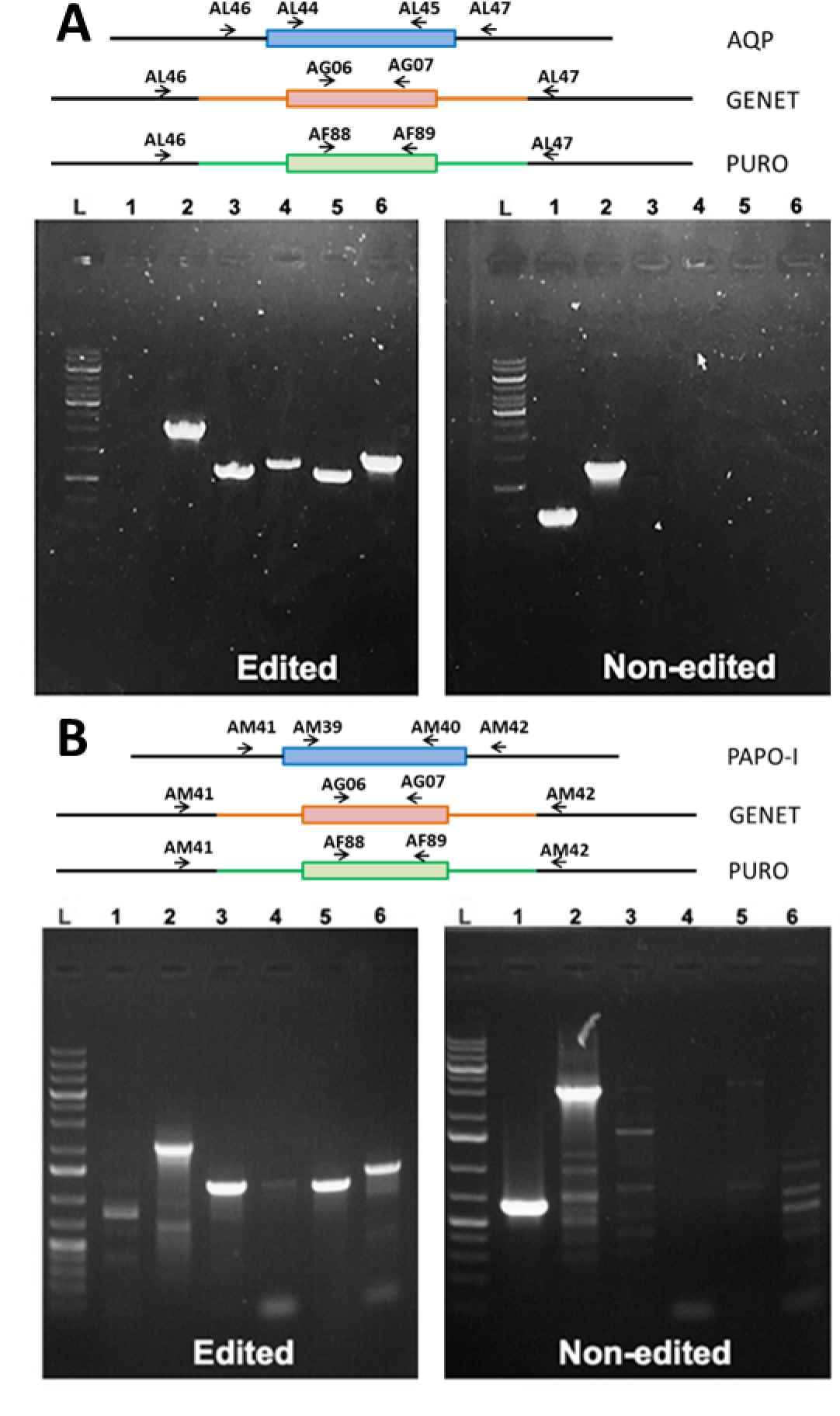
Validation of LdAQP KO and LdPAPO-I KO. Upper panels: schematic representation of each gene locus and primers (black arrows) used to confirm integration of the different drug resistant markers and loss of wild-type allele in the respective cell lines. Geneticin (GENET), Puromycin (PURO) and Neomycin (NEO). Wells: L is GeneRuler^TM^ 1kb Plus DNA ladder, wells 1 to 6 represent the different primer pairs used to check for edition. (A) LdAQP. 1: AL44/AL45, negative in edited and positive at 660 bp in non-edited cells, 2: AL46/AL47: Positive: 1239 bp (non-edited), 1959/2056 bp (edited – Geneticin/Puromycin), 3: AL46/AG07: 1137 bp (edited), 4: AG06/AL47:1268 bp (edited), 5: AL46/AF89: 1114 bp (edited), 6: AF88/AL47: 1409 bp (edited); [B] PAPO-I. 1: AM39/AM40, negative in edited and positive at 23 bp (non-edited), AM41/AM42: Positive: 2986 bp (non-edited), 2014/2111 bp (edited – Geneticin/Puromycin), AM41/AG07: 1196 bp (edited), AG06/AM42: 1263 bp (edited); AM41/AF89: 1173 bp (edited) and AF88/AM42: 1405 bp (edited).

For episomal expression (add-back and overexpression), LdAQP gene was cloned in a pTH6nGFPc vector (40) in a frame with a GFP tag and transfected into LD1S promastigotes. The quantitative expression of the AQP gene was checked and confirmed with qRT-PCR to confirm the overexpression of the AQP gene. qRT-PCR of the AQP presents a ∼7-fold increase compared to the wild-type LD1S cell (Fig. 3). The fluorescence due to the GFP tag also confirmed the presence of the AQP episomal expression (Fig. 4). GFP tag was used to determine the localisation of the protein in the cell. The localisation of AQP-GFP the localization is made up of a fine network and few more intense dots. An intense anterior dot was already noticed in Biyani et al. 2011 work (52). To better define the localisation observation, the GFP-tagged AQP proteins were co-localized with Mitotracker (Fig 4). Co-localization of the GFP and Mitotracker staining allowed us to determine that Ld AQP localizes to the mitochondrion. AQP KO parasites were obtained using a PCR-based CRISPR-Cas9 strategy (39). We validated the KO by the targeted integration of the two resistance genes

**Fig 3:**
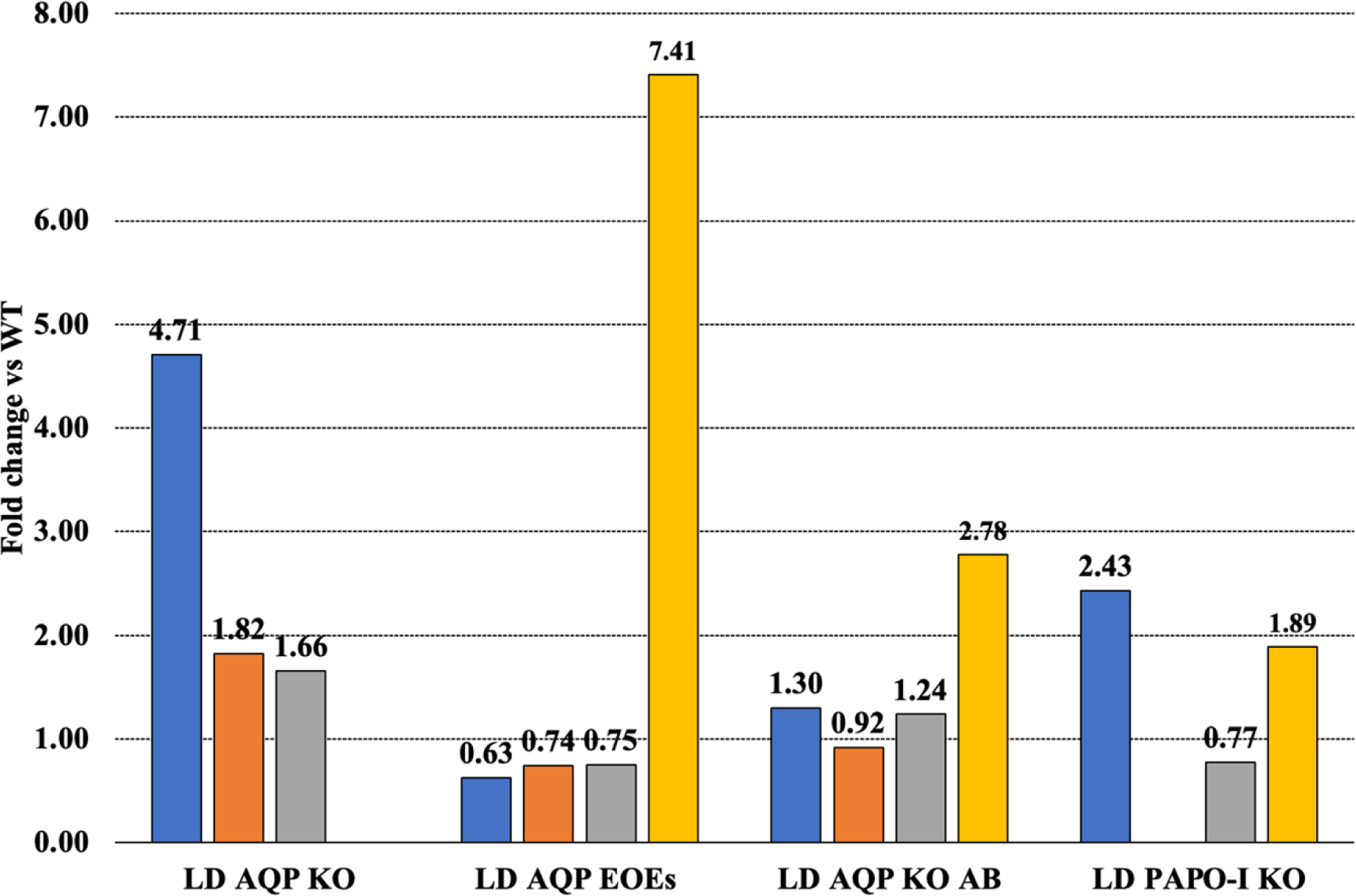
Quantitative expression mutants. Histograms show the fold change in gene expression when compared to the WT. Blue correspond to AAPO; orange to PAPO-I; grey to PAPO-II; and yellow to AQP.

**Fig. 4:**
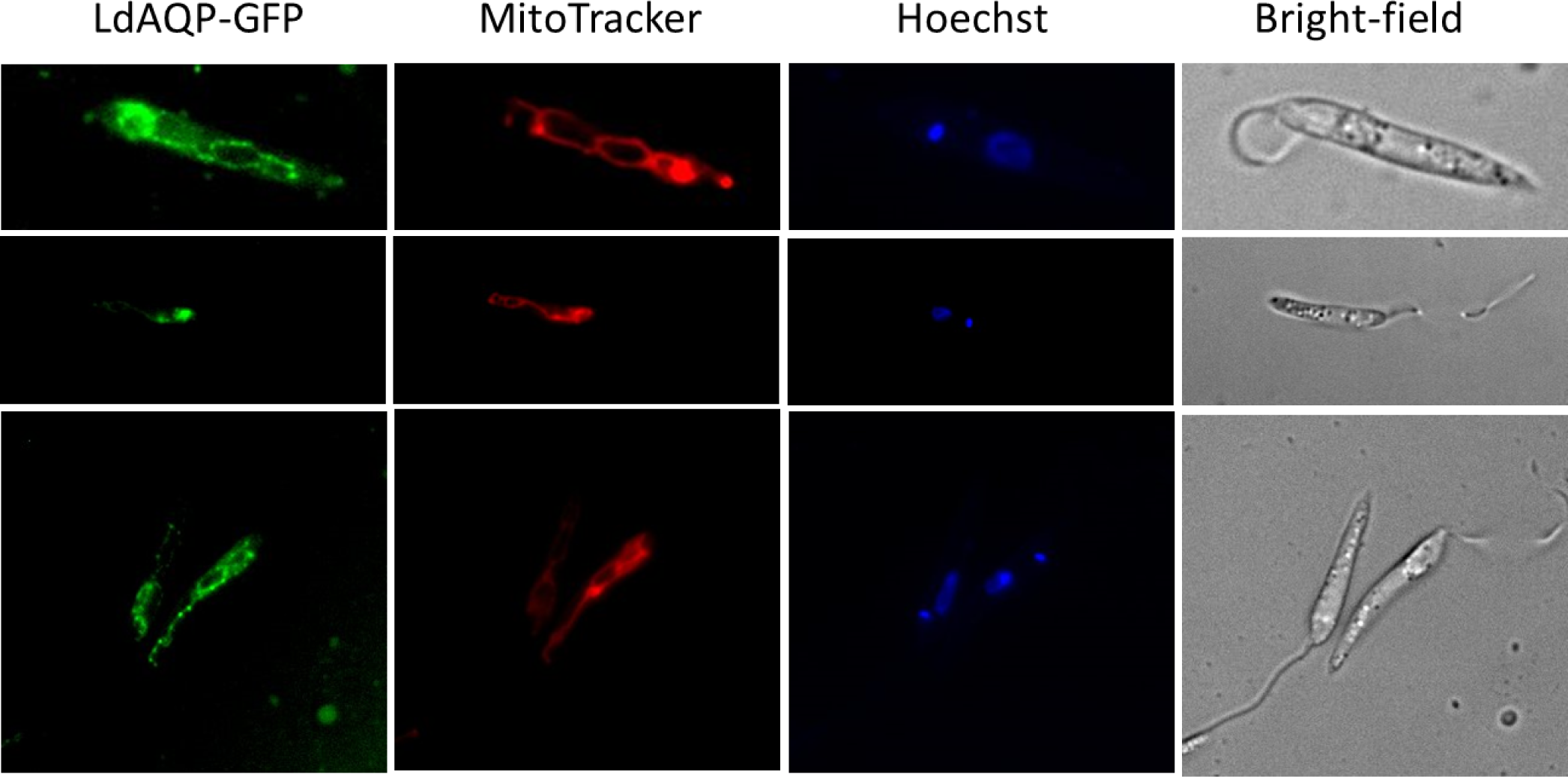
Localization of LdAQP. A fine network typical of the mitochondrion was obtained in LdAQP-GFP (green) and Mitotracker (red) staining. In some cells, as in the first row, a more intense staining is visible in the anterior part of the parasite, distinct to the flagellar pocket.

(Puromycin & Geneticin) and the non-amplification of the target gene by PCR (Fig. 2). This absence of gene-specific bands on the gel enabled us to conclude that we had obtained a pure KO strain after cloning. Deletion of AQP was also checked by the absence of expression of the AQP gene with qRT-PCR (Fig. 3). After that, we performed other qRT-PCRs to look into whether the absence of AQP modulate the expression of AAPO, PAPO-I, and PAPO-II developed in various AQP deleted mutant parasites. The expression level of AQP influenced the expression of all three genes. When AQP expression in KO parasites is zero, the expression of the other three increases from around 2-fold (PAPO-I and PAPO-II) to 5-fold for AAPO. Overexpression of AQP by a factor of 7 tended to reduce the expression of the other three genes. In the control complementation (in which a slight overexpression of AQP by a factor of 3 was observed), the expression of AAPO, PAPO-I and PAPO-II was unchanged compared with the WT (Fig. 3). These data favoured the hypothesis that these proteins form a complex, as suggested by the molecular docking studies. We then wanted to see if AQP was implicated in a higher sensitivity or resistance to an apoptotic stimulus. To do so, we used low doses of MLT. At 5 and 10 µM MLT, the TUNEL assay only detects ≤1% of apoptotic cells in WT and AQP KO cells (Fig. 5). However, in the AQP episomal overexpressing parasites, the percentage of apoptotic cells is increased to 11% and 24% in a dose-dependent manner (Fig. 5). Of note, there is a slight increase of apoptotic cells in the AQP add-back parasite to 4%, which is probably linked with the slight overexpression (∼3 fold) of AQP in that condition. These data suggested that AQP has a role in the apoptotic pathway in *Leishmania*.

**Fig. 5:**
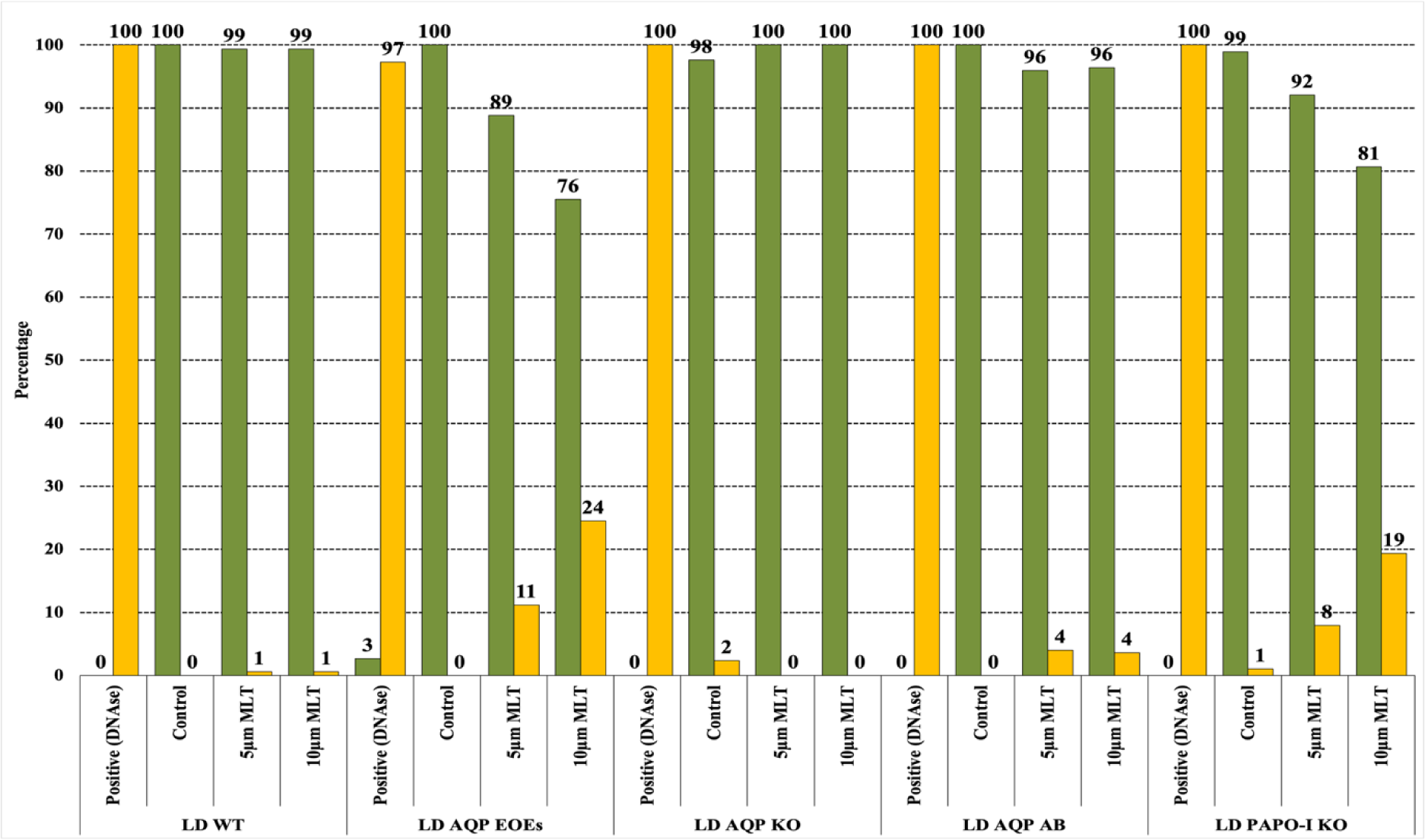
Exacerbated effect of MLT in mutant parasites: LD WT, LD AQP EOEs, LD AQP KO, LD AQP AB, and LD PAPO-I KO cells were treated with 5 and 10μM MLT. DNase treated cells were used as positive control. Apoptotic cells were detected using TUNEL assay. Histograms represents the percentage of Tunel positive cells (yellow) and Tunel negative cells (green).

#### Investigating the effects of identified apoptotic hit proteins on Leishmania apoptosis-like events

To assess the implications of our other identified apoptotic hit proteins, we employed a similar PCR-based CRISPR-Cas9 KO approach as for AQP. We attempted to knock out the other genes: AAPO, PAPO-I, and PAPO-II. A strategy similar to that used for AQP, employing specific primers in conventional PCR, was employed to verify the proper integration of the resistance marker and the complete absence of the target gene. We succeeded to obtain true PAPO-I KO mutant parasites. Here, the correct integration of the resistance marker and the complete absence of the PAPO-I was confirmed by PCR (Fig. 2B), which was further affirmed by the absence of expression by qRT-PCR. (Fig. 3). As in LdAQP KO parasites, we also observed a shift in the quantitative expression profile of AAPO, PAPO-II and AQP. AAPO and AQP were overexpressed around two-fold, and gene expression of PAPO-II was stable or decreased slightly by 0.77-fold of the value in the WT (Fig. 3). When we checked for sensitivity or resistance to apoptotic stimuli, PAPO-I KO was observed to be more sensitive to MLT. PAPO-I KO treated with MLT presented dose-dependent sensitivity with TUNEL-positive cells at 8% and 19%, respectively, for the 5 and 10 μM concentrations (Fig. 5). In two independent experiments, we attempted to knockout the AAPO gene without success. We also failed to obtain a true KO of PAPO-II, like for AAPO, the locus was editable, but despite cloning, a wild-type copy of the targeted gene persisted, despite the correct integration of two selective marker.

#### Analysing the quantitative expression of hit in MLT treated mutant cells

We examined the quantitative gene expression of AAPO, PAPO-I, PAPO-II, and AQP in MLT-treated mutant cells approximated to the wild type. Quantitative expression by qRT-PCR for genes was normalized again with cGAPDH. Expression profile (Fig. 6) presents a consistent decrease AQP expression in a dose-dependent manner in all mutants from 5 to 10 μM MLT treatment, while in LD AQP KO AB & LD PAPO-I KO mutants, the expression increased at first with 5 μM MLT treatment compared to a control cell and then decreased linearly from 5 to 10 μM MLT treatments in both the mutants. Meanwhile, AAPO, PAPO-I, and PAPO-II expression present a dull increase or decrease (insignificant) from control to 5 to 10 μM MLT treatments. Expression of AAPO remains unchanged in all the mutant control and 5 μM MLT-treated cells, with a slight increase (insignificant) in the case of 10 μM MLT treatment. PAPO-I expression decelerated slightly from control to 5 μM MLT treatment and remained unchanged in 10 μM MLT treatment. PAPO-II expression increases in 10 μM MLT treated LD AQP EOEs cells, while decreased and increased in a dose-dependent manner in LD AQP KO AB & LA PAPO-I KO, respectively.

**Fig. 6:**
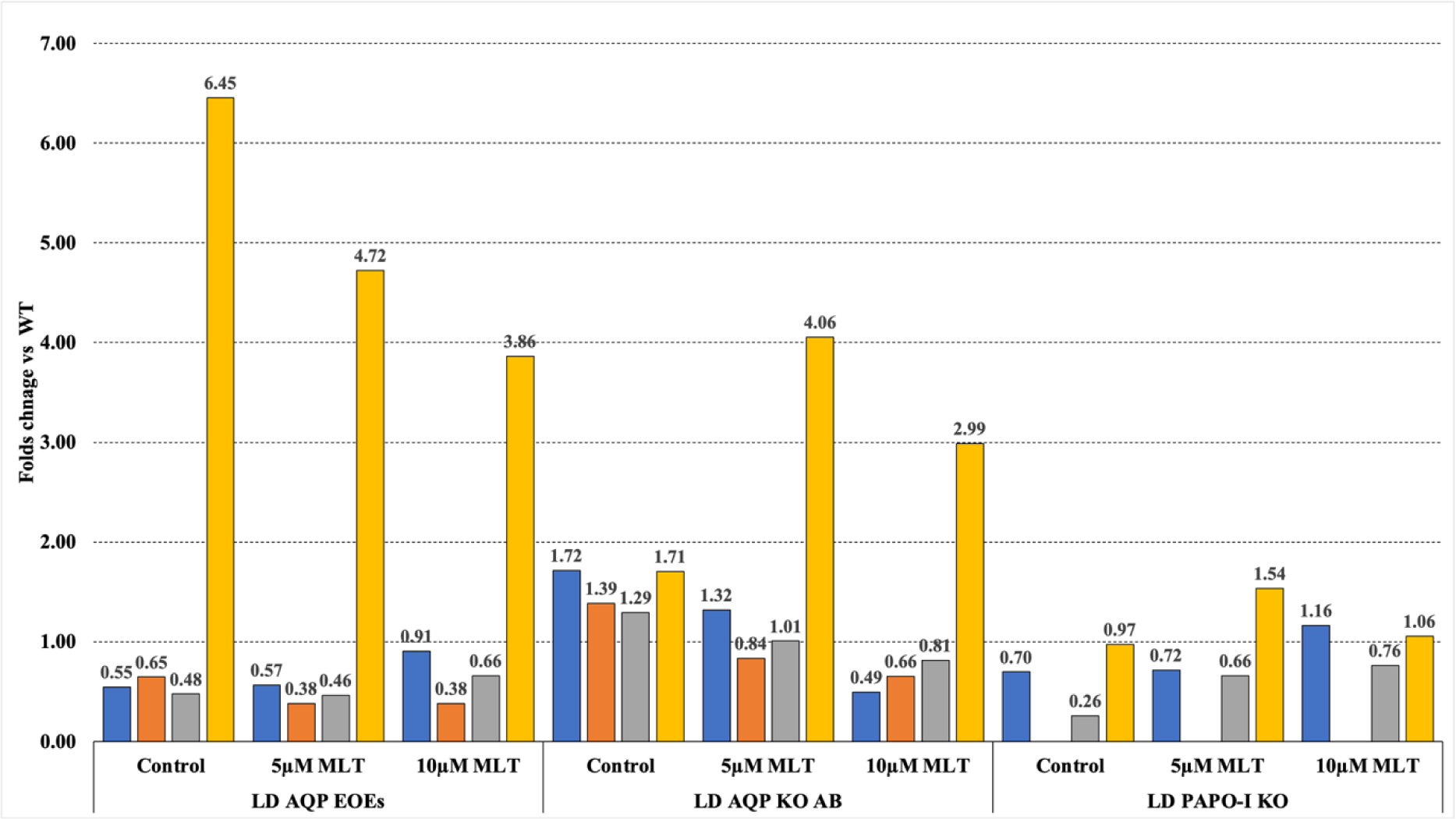
Quantitative expressions of the apoptotic genes in MLT treated cells in mutant parasites: Fold change in AAPO in blue, PAPO-I in orange, PAPO-II in grey and AQP in grey the mutants LD AQP EOEs, LD AQP KO AB, LD PAPO-I KO when treated with varying concentration of MLT in comparison to untreated cells.

## Discussion

Apoptosis in higher eukaryotes, such as mammals, is a well-studied process with extensive knowledge about its molecular mechanisms, regulation, and physiological significance in development, tissue homeostasis, and disease. In contrast, apoptosis in unicellular divergent eukaryotes, particularly in organisms like *L. donovani*, is relatively understudied and less understood. We searched for evolutionarily conserved proteins and validated their role in an apoptosis-like process in *L. donovani*.

### CRISPR-Cas9 gene edition in L. donovani

We successfully adapted the CRISPR-Cas9 PCR-based strategy developed in *L. Mexicana* (39) to *L. donovani.* Using this method, we succeeded to edit LdAQP and LdPAPO-I and to obtain null knock out strains. However, we attempted to knockout the AAPO gene without success, suggesting its essential role in *Leishmania* promastigotes. This finding aligns with previous research indicating the essentiality of its ortholog in *T. brucei* (Tb927.6.3870) in procyclics, as determined by high-throughput phenotyping using RNAi target sequencing (RIT-seq) (53). We also failed to obtain a true KO of PAPO-II, like for AAPO, the locus was editable, but despite cloning, a wild-type copy of the targeted gene persisted, despite the correct integration of two selective marker. This is likely attributed to the flexibility of the *Leishmania* genome and the presence of mosaic aneuploidy, which contribute to the difficulty in obtaining stable knockout mutants, even when multiple selective markers are employed (54–57). Several attempts to remove all copies of PAPO-II were unsuccessful, including a knock-in strategy. The situation is complicated by the presence of two copies of the gene at two loci, one on chromosome 2 and the other on 27, and the fact that the latter is not annotated on the LdBPK genome. Of note two copies are present in the genome of most sequenced *Leishmania* species (LdCL_020012400-t42_1 and LdCL_270033400-t42_1, LINF_020012400-T1 and LINF_270033830-T1, LmjF.02.0710:mRNA and LmjF.27.2630:mRNA) and also in *T. brucei* (Tb927.2.3030 and Tb927.2.5980). A knock-in strategy supposedly not dependent on genome misannotation also failed. This suggests that this duplicated gene is essential for *Leishmania* promastigote growth. In the high-throughput phenotyping using RNAi target sequencing (RIT-seq) data, Tb927.2.3030 and Tb927.2.5980, which have non-homologous nucleotide sequences, are not essential genes, but this could be because they are functionally redundant.

### Tentative model for apoptosis in L. donovani

Combining our *in silico* and experimental findings, our screened proteins AAPO, PAPO-I, PAPO-II & AQP are a few stakeholders of *leishmania*’s apoptosis or apoptosis-like event. We have demonstrated that AQP deletion makes the parasite resistant to MLT, and overexpression makes it sensitive to MLT, proposing it a pro-apoptotic protein. AQP being pro-apoptotic is logical because it possesses the BH3 domain. This BH3 domain is present and is responsible for binding mammalian pro-apoptotic pore former Bcl-2 proteins. (Fig. 4). The fine network observed with the GFP was highly suggestive of the single mitochondrion that characterize these parasites; the colocalzation with Mitotracker confirmed the mitochondrial localization of LdAQP. Additionally, in BLAST analysis, the highest score obtained for LdBPK_221270.1 (LdAQP) matches human Aquaporin 8, which is known to be a mitochondrial aquaporin, further supports the mitochondrial localization of LdAQP. Combining and comparing the TUNEL results and quantitative expression profile of MLT treatment on mutant cells, it is exciting to notice that AQP expression is decreased in a similar fashion with an increase in MLT concentration. Moreover, AQP and PAPO-I are making cells sensitive to MLT treatment. AQPs are a family of transmembrane water channel proteins required for water homeostasis. In addition to their primary function as water channels in mammals, they are essential for transporting glycerol, small solutes, and molecules, including carbon dioxide, metalloids, nitric oxide, ammonia, urea and various ions(58). Loss of cell volume or apoptotic shrinkage is an essential process in the cells entering the apoptotic pathway. AQPs, being influential in affecting the rate of water movement across the cell membrane, have been associated with the apoptotic volume decrease (AVD)(59) and subsequent apoptosis. Jablonski et al., in their study in ovarian granulosa, have shown the inhibiting AQPs blocked AVD and apoptotic events such as cell shrinkage, changes in the mitochondrial membrane potential, DNA degradation, and caspase-3 activation and, when overexpressed, showed loss of water permeability and higher apoptotic induction. Another study showed that overexpression of AQP confers resistance to apoptosis (60). These mixed findings from research suggest the possible role of AQPs in facilitating AVD necessary for apoptosis, and also, the overexpression of AQPs could make cells resistant to apoptosis. Another exciting and evident finding about aquaporins in *T. cruzi* is its presence in the acidocalcisomes and contractile vacuole, suggesting its role in the osmotic adaptions of parasites (61). Acidocalcisomes, in particular, are the compartments supposed to sequester the *L. donovani* metacaspases (62) in its acidic environment, which are synthesised in active form and released during the parasite apoptotic event. Our findings with AQP point to its role in *L. donovani* apoptosis, which is a big step forward in the molecular characterisation of apoptotic pathways in *Leishmania*. However, the relevance of AQP in osmoregulation, parasite homeostasis, and the function of the BH3 domain remains unknown.

According to our preliminary data on PAPO-I, it rather be considered as an anti-apoptotic protein, whereas our *in-silico* search based on mammalian pro-apoptotic Bcl-2 protein is supposed to be pro-apoptotic from. PAPO-I knockout cells were sensitive to MLT, and further complementation and localization of PAPO-I is required to conclude it as pro or anti-apoptotic proteins. Variation in the quantitative expression profile of all the identified proteins in mutants in the presence or absence of MLT as apoptotic stimuli suggests they somehow interact and are important during *Leishmania*’s apoptosis or apoptosis-like events. However, more experiments are needed to describe the mutant parasite’s cellular defects finely and in-vitro interacting partners.

### A Novel Therapeutic Target Discovery Approach

The chemotherapeutic approaches employed for treating leishmaniasis have been associated with numerous side effects and the development of resistance. Immunotherapy and immunochemotherapy are newer therapeutic approaches gaining popularity, but they are still in their infancy. Understanding pathogenesis and pathogen’s metabolic pathways has led to the creation of several medications that target distinct biochemical pathways. We found encouraging findings for apoptotic protein in *L. donovani*, in this study and a potential apoptotic pathway. In particular, our data suggested that AAPO and the duplicated PAPO-I gene were essential, making them targets of choice for therapeutic interventions. It was accomplished using orthologous gene screening in *L. donovani versus* key mammalian Bcl-2 apoptotic proteins, molecular docking investigated their interaction, and molecular dynamic simulation studies did interaction validation. This research finding clearly suggest unequivocally that the protein-protein complex combination III is a potentially crucial phase in the apoptotic process of *L. donovani*. This is a first-of-its-kind work that identifies apoptotic proteins and pathways in *L. donovani* and presents a novel therapeutic target. We feel that further confirmation of the suggested protein-protein interaction would need experimental validation. Furthermore, the technique developed here yielded remarkable findings and may be used simultaneously to predict and uncover apoptotic protein orthologs in other diseases, as well as to foster multidisciplinary partnerships of expertise from experimental and computational scientists.

## Acknowledgement

KK is thankful to the Central University of Rajasthan for offering a fellowship and EMBO for awarding a Scientific Exchange Grant to visit the YS lab at Université de Montpellier. VKP is thankful to the Central University of Rajasthan for the computing facility and infrastructure required to complete the project. VKP is thankful for IRD support for this collaborative research activity and funding from the Institute of Eminence, University of Delhi, and the Indian Council of Medical Research (IIRPIG-2023-0000873). KK & VKP acknowledge YS for hosting the experimental validation. VKP & YS are thankful for Labex-CeMEB doctoral mobility under the Université de Montpellier I-SITE excellence program for subsistence. KK is also thankful to Maude Lévêque for the insightful suggestion during the project. YS received funding by the Agence Nationale de la Recherche within the frame of the “Investissements d’avenir” program (ANR-11-LABX-0024-01 “PARAFRAP”).

## Authors contribution

Ketan Kumar: Data curation, Conceptualization, Visualization, Methodology, Software, Experimental work, Writing-original draft, Data interpretation, Validation. Lucien crobu: Data curation, Conceptualization, Visualization, Methodology, Experimental work. Yvon Sterkers: Conceptualization, Experimental work, Visualization, Validation, Resources, Investigation, Data interpretation, Supervision, Writing-review & editing. Vijay Kumar Prajapati: Conceptualization, Validation, Investigation, Supervision, Writing-review & editing.

---

**Supplementary Fig. 1:**
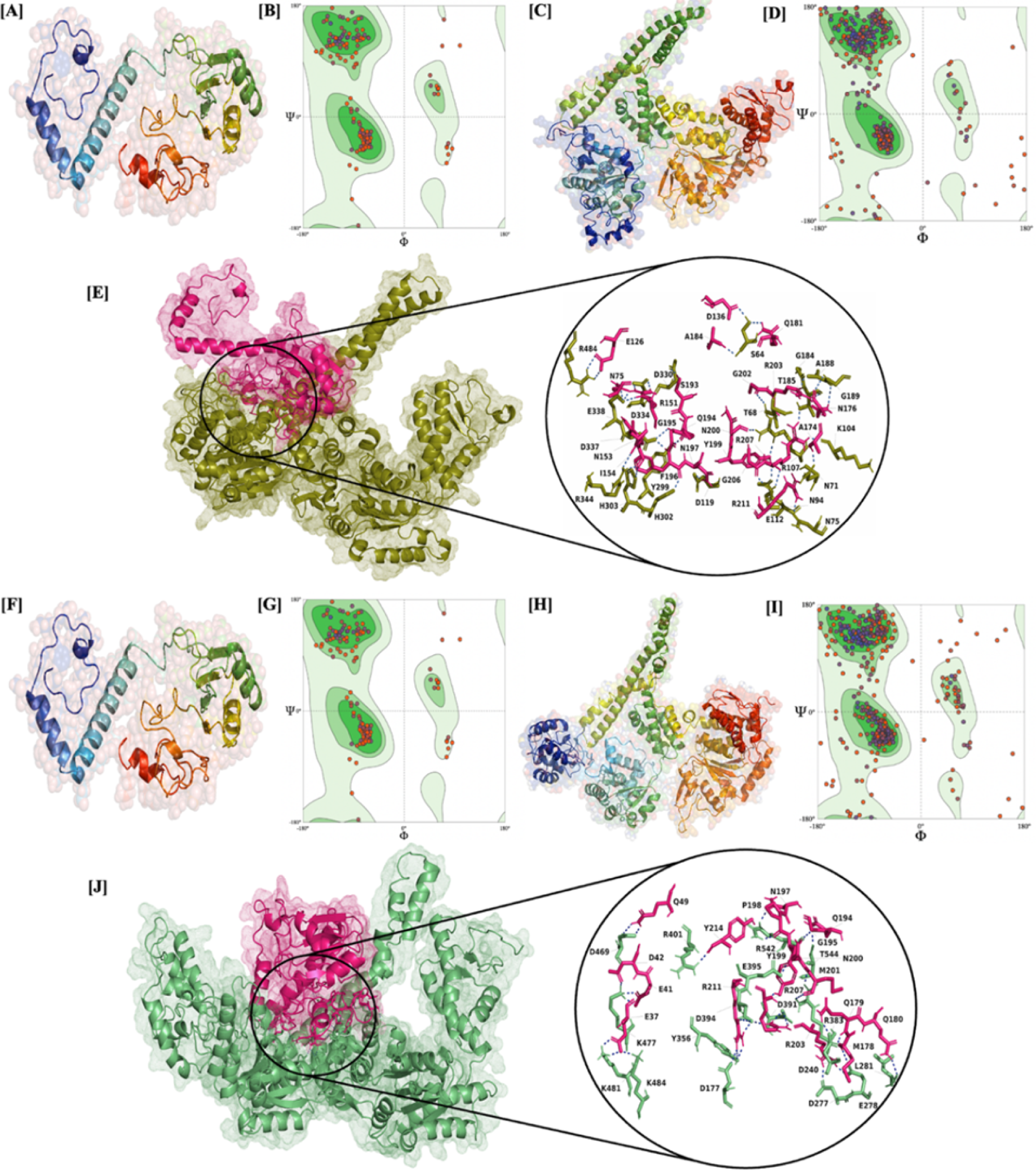
In silico structure and binding prediction of different orthologs. [A] Homology structure modelled by Phyre2 for AAPO; [B] Ramachandran plot for ϕ and ψ angles for the validation of homology predicted AAPO structure; [C] Homology structure modelled by Phyre2 for PAPO-I; [D] Ramachandran plot for ϕ and ψ angles for the validation of homology predicted PAPO-I structure. [E] Docking prediction of AAPO in pink and PAPO-I in green with their interacting residue in the inset image. [F] Homology structure modelled by Phyre2 for AAPO; [G] Ramachandran plot for ϕ and ψ angles for the validation of homology predicted AAPO structure; [H] Homology structure modelled by Phyre2 for PAPO-II; [I] Ramachandran plot for ϕ and ψ angles for the validation of homology predicted PAPO-II structure. [J] Docking prediction of AAPO in pink and PAPO-II in olive green with their interacting residue in the inset image.

**Supplementary Fig. 2:**
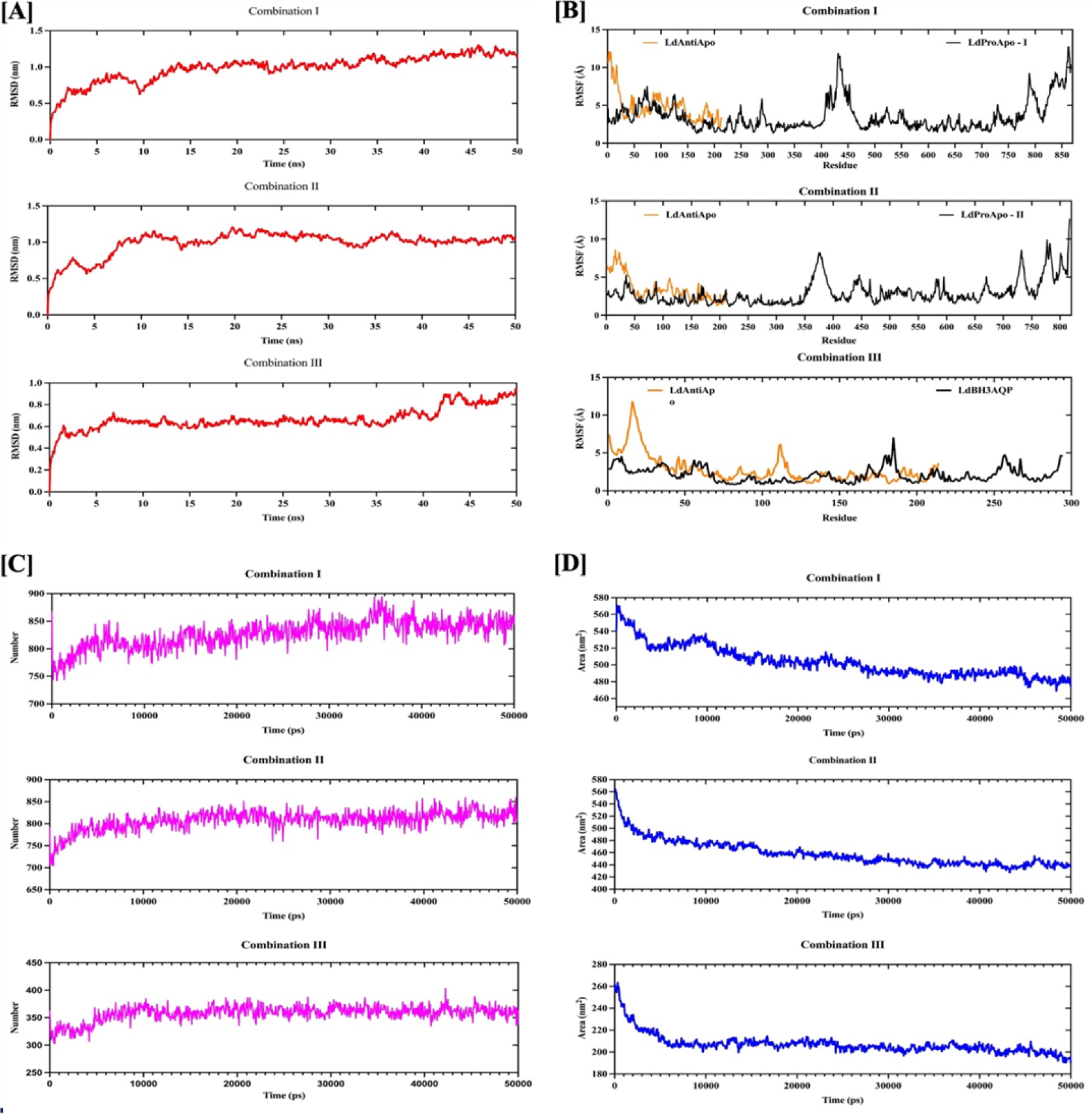
Trajectory graph evaluating dynamic behaviour and conformation consistency of protein-protein complex over time. The graph illustrates the RMSD for Cα atom coordinates coloured in red in Combinations I, II, and III in nanometres (nm) versus the initial structure as a function of time in nanoseconds (ns) [A]. The graph illustrates the RMSF of Cα atom coordinates coloured in Black for Pro-apoptotic proteins and Orange for Anti-apoptotic proteins of Combinations I, II, and III as a function of residue index versus in nanometres (nm) [B]. The graph illustrates the Radius of gyration (Rg) coloured in purple for Combinations I, II and III in nanometres (nm) as a function of time in nanoseconds (ns) [C]. The graph illustrates the SASA coloured in blue for Combinations I, II and III in nanometres square (nm2) as a function of time in nanoseconds (ns) [D].

**Supplementary Table 1:**
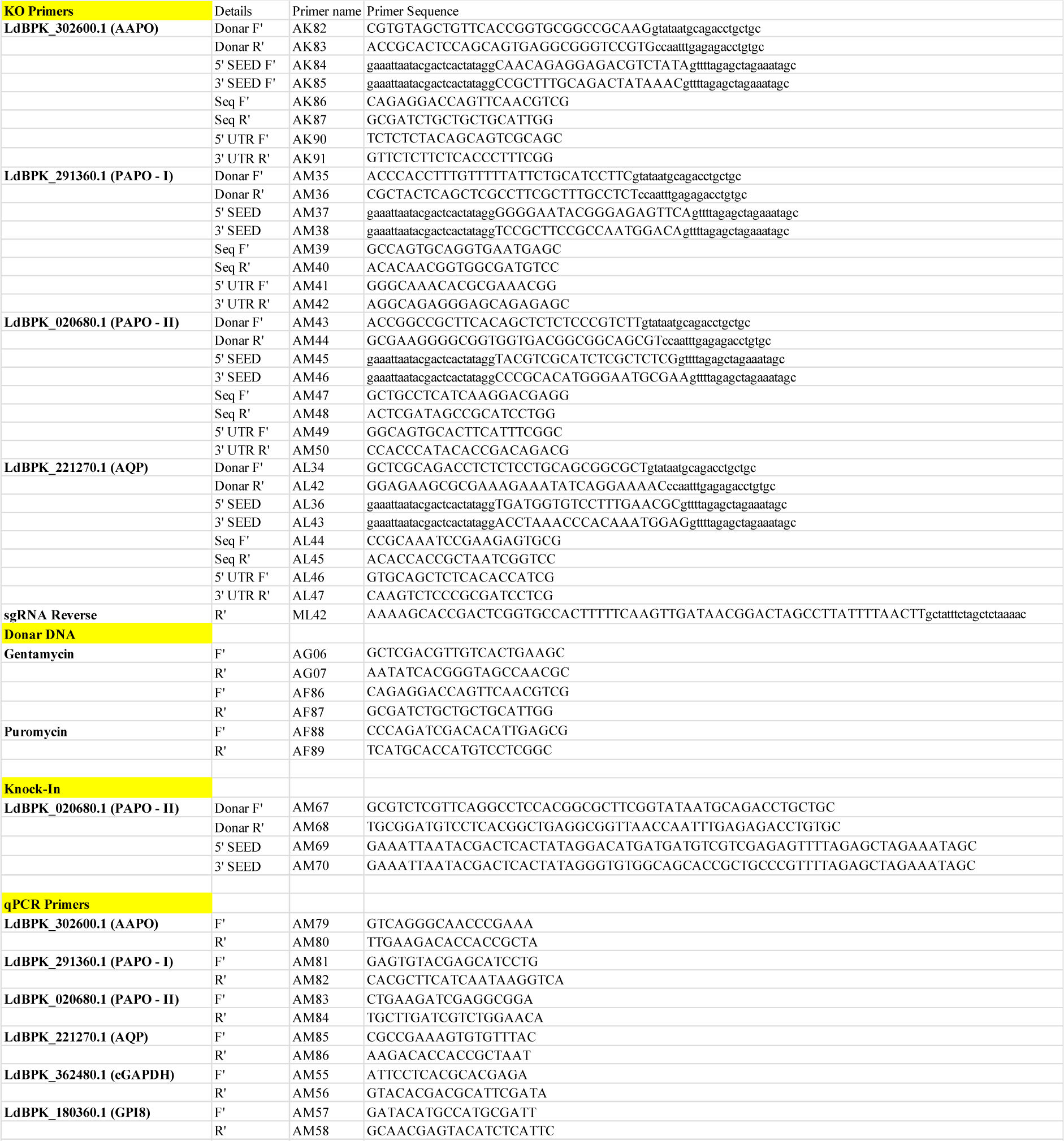
Primers included in the study

